# Sex Differences in the Effects of Controlled Circadian Dysregulation on Gut Microbiome and Intestinal Barrier Architecture

**DOI:** 10.64898/2026.01.24.701506

**Authors:** Ella Barnum, Jonathan L. Turck, Karienn A. Souza, Kathiresh Kumar Mani, Rachel Pilla, Amutha Selvamani, Farida Sohrabji, David J. Earnest

## Abstract

Disturbances of 24-hour or circadian rhythms imposed by everyday irregular work and/or social schedules have been linked to vascular disease, including ischemic stroke. Using an established shift work-like paradigm and preclinical model for ischemic stroke, we have shown that environment-induced circadian dysregulation exacerbates stroke outcomes differentially to a greater extent in male than female rats. Because more severe stroke outcomes and circadian rhythm disturbances have been linked to gut pathophysiology, present study examined the effects of chronic LD cycle shifting on gut cytoarchitecture, microbiota composition, metabolites, and gut-derived inflammatory mediators. Adult (5-7mo) rats were divided into 2 groups and exposed for 50d to a fixed or shifted (lights-on advanced by 12hr/5d) LD 12:12 cycle. Circadian entrainment of activity rhythms was stable in all rats on the fixed LD 12:12 cycle but was severely disrupted during exposure to shifted LD cycles. Significant changes in the composition of the gut microbiome including reduced alpha diversity, shifts in beta diversity and correlations between the abundance of beneficial gut bacteria and stroke survival were observed in male but not female rats exposed to shifted LD cycles relative to fixed LD controls. This effect of circadian dysregulation on gut microbiota was accompanied by evidence of pathologic gut morphology (i.e., shorter and blunted villi, crypt hyperplasia disruption of tight junction proteins and gut barrier integrity), elevated serum endotoxin concentrations, decreased levels of the short-chain fatty acid (SCFA) butyrate, and increased circulating levels of the inflammatory cytokine IL-17A in shifted LD male rats. These results suggest that alterations in gut morphology, microbiota and metabolites may contribute to sex differences in the effects of shift work-related circadian dysregulation on ischemic stroke outcomes.

## INTRODUCTION

The generation and photoentrainment of mammalian circadian rhythms is mediated by a hierarchical organization of multiple, cell-autonomous clocks. The suprachiasmatic nucleus (SCN) of the anterior hypothalamus functions as a master pacemaker whereas peripheral clocks located in other brain regions and tissues provide for the local coordination of tissue- or cell-specific processes so as to occur at the “right time” of day or night relative to each other and to environmental time cues (Bell-Pedersen et al., 2015). Dysregulation of circadian clocks throughout the body and corresponding misalignment of local tissue- and cell-specific rhythms has been commonly linked to shift work, chronic jet lag, and workplace or social influences that commonly impose highly irregular schedules on our sleep-wake patterns, mealtimes, and other body processes. In this regard, it is noteworthy that about 20-25% of the US population engage in shift work at non-traditional times during evenings, nights, or rotating shifts, and the resultant dysregulation/misalignment of circadian rhythms has been identified as a contributing element in vascular disease or related risk factors (e.g., diabetes, obesity, inflammation) in general and stroke in particular (Marcheva et al., 2010, Mosendane et al., 2008, Reilly et al., 2007, Scott et al.,2008, Tuchsen et al., 2006, Turek et al., 2005, Westgate et al.,2008, Torquati et al., 2018, Karlsson et al., 2005).

Because shift work is associated with other major risk factors for vascular disease, such as smoking, poor diet and lower socioeconomic status, these epidemiological studies provide limited opportunity to resolve the extent to which circadian rhythm dysregulation by shift work schedules exclusively contributes to cardiovascular and stroke pathology. Our recent studies with animal models have provided an important complement to epidemiological observations by using chronic shifts of the light-dark (LD) cycle to determine whether shift work-like modulation of circadian rhythms alone exacerbates the extent of stroke-induced brain injury and functional impairment. In adult (5-7mo) rats, we found that circadian rhythm dysregulation in response to recurrent advances (12hr/5d) of the LD cycle amplifies sex differences in pathological outcomes of ischemic stroke immediately following exposure to shifted LD cycles (Earnest et al. 2016). Similar to reported epidemiological findings (Karlsson et al., 2005), circadian dysregulation after exposure to shifted LD cycles was acutely marked by high rates of stroke-induced mortality in males (70%), but not females (20%). At present, little is known about the mechanisms by which shift work-induced circadian dysregulation amplifies stroke severity and how this variable interacts with other non-modifiable risk factors such as biological sex to modulate the pathological effects of stroke.

Our studies and other published reports have implicated two broad and overlapping mechanisms involving inflammation and gut-brain communication in mediating the effects of circadian dysregulation on stroke outcomes. Stroke is an inflammatory disease and post-stroke disruption of the blood brain barrier as well as activation of systemic and brain resident immune cells are key factors in the pathophysiology of stroke. Consistent with the contributions of inflammation and immune cell activation to stroke-induced brain injury and functional deficits, circadian rhythm dysregulation has been shown to promote proinflammatory responses of the innate immune system, leading to a persistent inflammatory condition (Castanon-Cervantes et al., 2010, Xu et al., 2014, Kim et al., 2018). The role of inflammation and the immune system in mediating the pathological effects of circadian dysregulation on stroke is indirectly supported by our finding that global disruption of circadian rhythms in mice with targeted clock gene mutations enhances activation of proinflammatory M1 macrophages and their inflammatory responses, thereby promoting tissue inflammation (Xu et al., 2014). Over the past decade, increasing evidence suggests that interactions between the gut and brain play an important role in both normal and pathologic function (Dinan and Cryan, 2017). Importantly, both stroke and the disruption of circadian rhythms have been shown to alter gut barrier integrity, microbiota composition and metabolites (Stanley et al., 2016, Deaver et al., 2018). Because the gut houses large cohorts of immune cells and gut metabolites that modulate inflammation, it is possible that circadian dysregulation may exacerbate ischemic stroke severity by promoting gut pathophysiology and consequently an inflammatory condition. Using an established light-dark (LD) cycle shifting paradigm, the objective of the present study was to investigate whether the exacerbation of stroke pathology in response to controlled circadian dysregulation is linked to alterations in gut barrier integrity, microbiota and beneficial metabolites, and elevations in circulating levels of inflammatory mediators.

## MATERIAL AND METHODS

### Animals

Adult (5–7-month-old) female and male Sprague Dawley rats were purchased from Harlan Laboratories and maintained in the AAALAC-accredited vivarium at the Texas A&M University Health Science Center under controlled temperature (22–25°C) and lighting (LD 12:12) conditions with food (standard rat chow) and water available *ad libitum*. All animals were housed individually in cages equipped with running wheels to provide for continuous analysis of wheel-running activity. All animal experiments were performed in accordance with the National Institutes of Health Guide for the Care and Use of Laboratory Animals. Animal procedures used in this study were conducted in compliance with Animal Use Protocol as reviewed and approved by the Institutional Animal Care and Use Committee at Texas A&M University.

To analyze the effects of circadian dysregulation, experiments used a chronic light-dark (LD) cycle shift paradigm that has been shown to be effective in desynchronizing circadian rhythms and in inducing pro-inflammatory responses of immune cells, leading to a persistent inflammatory condition (Castanon-Cervantes et al., 2010, Xu et al., 2014, Kim et al., 2018). After baseline acclimation under standard LD 12:12 conditions (lights-on at 0600hr; light intensity = 110-170 lux at 500-580nm) for about 2 weeks, animals were randomly divided into 2 groups and exposed for 50 days to either the same “fixed” LD 12:12 cycle or to a “shifted” LD 12:12 cycle. During exposure to the shifted LD paradigm, lights-on was advanced by 12hr (at 1800hr) every 5 days and these shifts in the LD cycle were repeated for 5 full cycles. At the conclusion of this treatment period, both groups were exposed to the same standard LD 12:12 schedule (lights-on at 0600hr).

### Analysis of wheel-running activity

To confirm the long-term effects of circadian dysregulation on the rhythm of wheel-running behavior, all fixed and shifted LD rats were housed individually in cages equipped with running wheels. Wheel-running activity was continuously recorded, stored in 10-minute bins, graphically depicted in actograms, and analyzed using ClockLab data collection and analysis software (ActiMetrics, Evanston, IL). Entrainment and qualitative parameters of the activity rhythm were analyzed as described previously (Earnest et al. 2016).

### DNA Isolation, Sequencing, and Bioinformatics

Analysis of the gut microbiome and metabolites in the circulation was performed on male (@n=16) and female (@n=16) rats (cohort 1) exposed to the LD cycle shift paradigm and subjected to ischemic stroke surgery in our original study demonstrating that circadian dysregulation exacerbates sex differences in ischemic stroke outcomes (Earnest et al. 2016). In this cohort, fecal and saphenous blood samples (pre- and post-treatment) were collected around mid-day of the LD 12:12 cycle (1100-1200hr) immediately before (baseline) and ≈4-5 days after the conclusion of experimental LD cycle manipulations when both treatment groups were exposed to the same LD 12:12 schedule (**Fig. 1**). Then all animals were subjected to ischemic stroke surgery at the same relative time during the circadian cycle (i.e., inactive phase; ≈ ZT 2-8) and stroke outcomes were assessed for 5 days. The fecal samples were frozen immediately after collection, and an aliquot of 100 mg (wet weight) of each fecal sample was used for DNA extraction using a MoBio Power soil isolation kit (MoBio Laboratories, USA) following the manufacturer’s instructions. A single-step 30-cycle PCR using the HotStarTaq Plus Master Mix Kit (Qiagen) was performed using primers 515F (5′-GTGYCAGCMGCCGCGGTAA -3′; Parada et al., 2016) to 806R (5′-GGACTACNVGGGTWTCTAAT -3’; Apprill et al., 2015) under the following conditions: 94°C for 3 minutes, followed by 28 cycles (5 cycles used on PCR products) of 94°C for 30 seconds, 53°C for 40 seconds and 72°C for 1 minute, after which a final elongation step at 72°C for 5 minutes was performed. Illumina sequencing of V4 region of the bacterial 16S rRNA genes was performed at the MR DNA laboratory (www.mrdnalab.com, Shallowater, Texas). The sequence data presented in this study are deposited in the National Center for Biotechnology Information (NCBI) Short Read Archive (SRA) database (Accession Number: PRJNA1334100).

**Figure 1:**
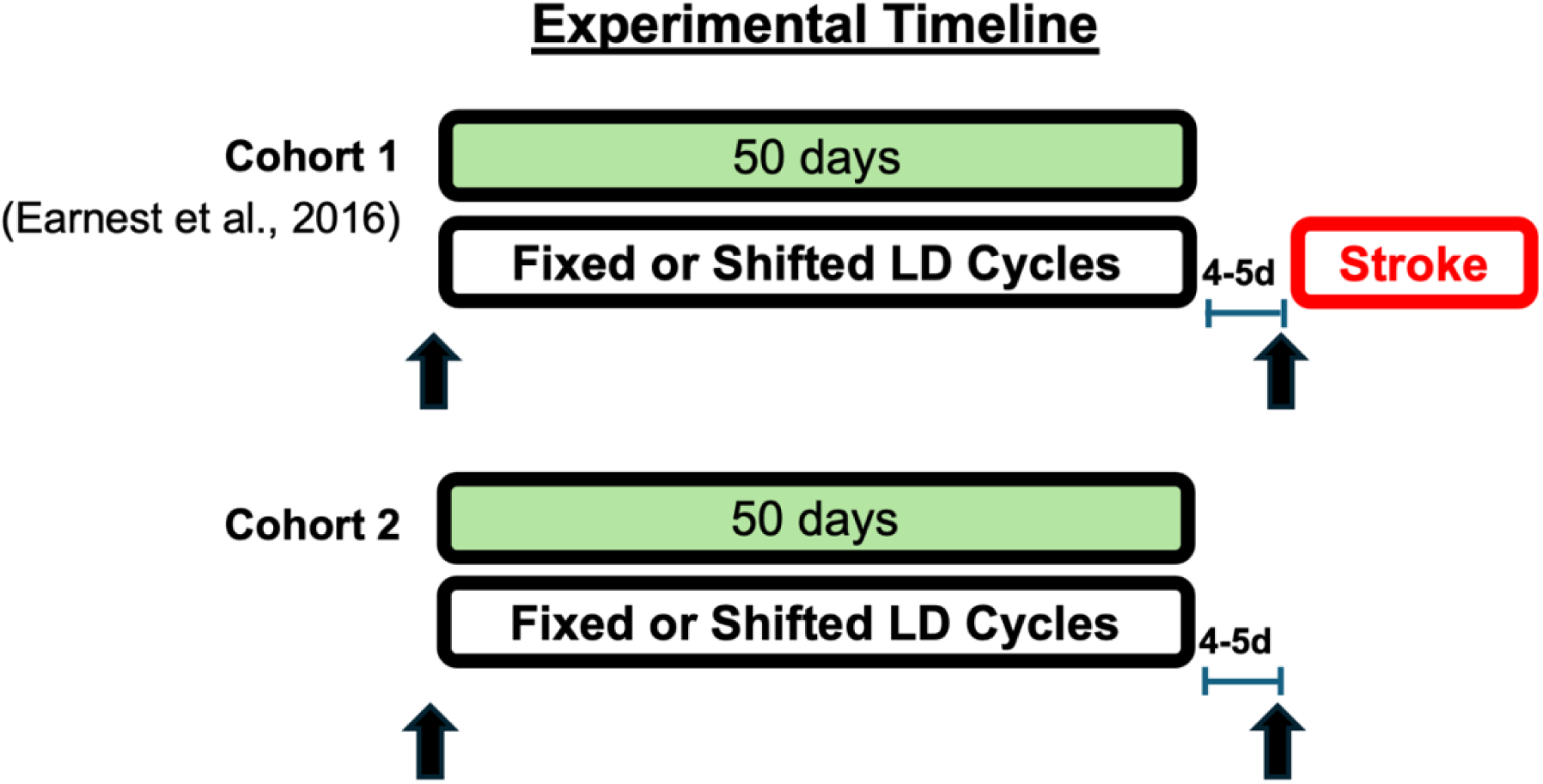
Experimental timeline. At 5-7 months of age, male and female rats were exposed for 50 days to fixed or shifted (12hr advance/5d) LD 12:12 cycles. Cohort 1 consists of male (@n=16) and female (@n=16) rats from our study examining the effects of circadian dysregulation on ischemic stroke outcomes (Earnest et al. 2016). In this cohort, fecal and saphenous blood samples (pre-and post-treatment) were collected around mid-day of the LD 12:12 cycle (1100-1200hr) immediately before (baseline) and ≈4-5 days after the conclusion of experimental LD cycle manipulations when both treatment groups were exposed to the same LD 12:12 schedule. Then all animals were subjected to ischemic stroke surgery at the same relative time during the circadian cycle (i.e., inactive phase; ≈ ZT 2-8) and stroke outcomes were assessed for 5 days. Cohort 2 represents separate groups of fixed and shifted LD rats (@n=8 males and n=8 females) to analyze gut morphology and barrier integrity. The timing of blood and/or tissue sample collection (pre- and post-treatment) is indicated by the arrows (Cohort 1: fecal samples and blood; Cohort 2: blood and portion of distal ileum).

The demultiplexed sequences were imported into QIIME2 (v. 2024.10) for analysis. The DADA2 denoising procedure (Callahan et al., 2016) was used to remove chimeric sequences and identify amplicon sequence variants (ASVs). To assign taxonomy, SILVA release 138 was used with the VSEARCH algorithm (Rognes et al., 2016). To determine phylogenetic relationships, sequences were aligned using MAFFT and a phylogenetic tree was constructed with FastTree (Price et al., 2010). Downstream analysis was performed in QIIME2 (v. 2024.10) and R (v. 4.5.0). A complete list of used R packages can be found in S1. The samples were then rarefied at the lowest per sample number of reads, 88,120 for the male dataset and 57,780 for the female dataset, for even diversity analysis depth. Alpha diversity was calculated using the observed features, and Shannon metrics to estimate community richness and evenness. A Shapiro-Wilk test was performed on the calculated alpha diversity metrics to assess normality. Both metrics were found to violate normality, thus a generalized linear mixed-effects models (GLMM; Gamma family, log link) were used. For all models the fixed effects were circadian condition, timepoint, and their interaction, while a random effect for animal (Animal_ID) was included to account for repeated measures (i.e., lm (alpha_diversity ∼ Condition * Timepoint + (1 | Animal_ID)). Estimated marginal means (emmeans) were then calculated for the condition and timepoint interaction. Pairwise post-hoc comparisons were calculated, and the resulting p-values were adjusted for multiple comparisons with the Tukey method. Beta diversity was calculated using weighted UniFrac dissimilarity to estimate differences in overall community structure. A permutational multivariate analysis of variance (PERMANOVA) model was used to assess significance using the vegan package (Anderson, 2001). The PERMANOVA model tested the effects of circadian condition, timepoint, and their interaction. To account for repeated measures, permutations were constrained with a blocking factor of the Animal ID across 999 permutations. Pairwise comparisons were then performed between all circadian condition and timepoint pairs. The resulting p-values were then corrected for multiple comparisons using the Holm-Bonferroni method. Additionally, intra-individual weighted UniFrac distance was extracted from pre- and post-pairs to assess individual change in beta diversity. Resulting distances between LD conditions were compared using a Wilcoxon rank-sum test.

The MaAsLin3 (v. 1.0.0) was used to identify differentially abundant and prevalent taxa at the genus level (Nickols et al., 2024). Only genera present in at least 10% of samples at a relative abundance of 0.01% were considered for differential abundance analysis. The MaAsLin3 linear mixed-effects models were fitted to fixed and shifted LD pre- and post-pairs with Animal ID included as a random effect, to account for repeated measures. To adjust for multiple comparisons p-values were corrected within MaAsLin3 using the Benjamini-Hochberg procedure. Following correction, joint q-values < 0.05 were considered significant for all above tests. Visualizations were produced in R with the packages ggplot2 (v. 3.5.2), phyloseq (v. 1.52.0), qiime2R (v. 0.99.6), ggprism (1.0.6), and ggsci (v. 3.2.0). Pearson’s correlation coefficients were determined to analyze the relationship between the abundance of specific bacterial taxa in fecal samples and stroke survival as gauged by the number of days (1-5) that animals survived after ischemic stroke.

Raw sequencing data have been deposited in the NCBI Sequence Read Archive (SRA) and are publicly accessible under the accession number PRJNA1334100. Code used for downstream analyses of 16S rRNA gene sequencing data is available on GitHub at https://github.com/JonathanTurck02/CD_gut_analysis.

### Short chain fatty acid analysis

Short-chain fatty acids (SCFAs) were analyzed in serum samples from animals in cohort 1 obtained immediately before and after exposure to experimental lighting conditions. Serum samples were aliquoted (50ul) and extracted with a methanol:chloroform:water-based extraction method. Samples were spiked with 0.1 mM d7 butyric acid as an internal standard. SCFAs (butyric acid, isobutyric acid, acetic acid, valeric acid, isovaleric acid, propionic acid) were detected and quantified on a gas chromatography triple quadrupole mass spectrometer (TSQ EVO 8000, Thermo Scientific, Waltham, MA) at the Texas A&M University Integrated Metabolomics Analysis Core.

### Gut histology, Immunohistochemistry and ELISA assays

Analysis of changes in gut morphology and in circulating levels of LPS and inflammatory cytokines required the collection of gut tissue and blood samples from an independent cohort that was not subjected to ischemic stroke because stroke itself: 1) is an inflammatory disease involving the activation of microglia, recruitment of peripheral inflammatory cells and secretion of inflammatory mediators (e.g., LPS, IL-17); 2) alters gut barrier integrity (Durgan et al., 2019; El-Hakim et al., 2021; Honarpisheh, Bryan and McCullough, 2022); and 3) serum volumes from saphenous blood samples obtained from rats in cohort 1 were insufficient for assaying circulating levels of inflammatory mediators. For analysis of the effects of circadian dysregulation on gut morphology and barrier integrity, a separate cohort of rats (**cohort 2**) were housed individually in cages equipped with running wheels and divided at ∼5mo of age into two treatment groups exposed to fixed or shifted (@n=8 males and n=8 females) LD cycles as described above (**Fig. 1**). Immediately before experimental LD cycle manipulations, saphenous blood samples were collected from all animals around mid-day of the LD 12:12 cycle (1100-1200hr). Following exposure to experimental lighting conditions (post-treatment), terminal blood samples and a portion of the distal ileum were collected at the same time during LD 12:12 cycle (1100-1200hr).

After dissection, the distal ileum was post-fixed, and embedded in Cryo-OCT compound (Leica Microsystems, Buffalo Grove, IL, USA). Embedded tissue blocks were stored at -80° until sectioning. Cryosections (10μm) were collected on glass slides and stained for hematoxylin and eosin (H&E) as previously described (Kumar et al., 2017). The sections were visualized and photographed with the FSX100 Cell Imaging System (Olympus) at × 10 magnification. To assess structural changes in gut morphology, villus width as well as the length of the villus and crypt layers were measured from these images using the ImageJ software. Based on previous methods (El-Hakim et al., 2021), villus dimensions were measured from the base at the crypt to its tip for villus length and at its widest point for villus width. The length of the crypt layer was measured from the base of the villus to the submucosal layer and muscularis externa. These measurements were assessed for every intact villus (10-15) on three sections per animal and reported as a ratio of villus length/width and villus length/crypt length.

Immunofluorescence for Zonula Occludens (ZO-1) was performed as previously described (Kumar et al., 2017). Cryosections (10μm) were collected on glass slides and incubated in blocking buffer (5% bovine serum albumin (BSA), 0.1% Triton X-100 in PBS, pH 7.4) for 1 h at room temperature. The sections were then incubated overnight at 4^0^C with primary antibodies to ZO-1 (rabbit polyclonal; Invitrogen, #61-7300) at 1:1000 and villin (mouse monoclonal; Santa Cruz, #58897) at 1:250 in PBS with 2.5% BSA. After rinses (3X) with PBS containing 0.5% Tween, slides were incubated with secondary antibodies (ZO-1: goat anti-rabbit IgG [H+L] Alexa Fluor 555 [ThermoFisher, Catalog #31210]; villin: goat anti-mouse IgG FITC [ThermoFisher, Catalog #A16085]) at 1:2000 dilution for 1 h at room temperature. The sections were then washed three times in PBS and coverslipped with mounting media containing the nuclear dye DAPI (Fluoroshield, Abcam). The sections were visualized and photographed on the FV12-IX83 confocal microscope. Subjective analysis of the distribution of ZO-1 immunofluorescence along brush border of intestinal villi was independently conducted by two investigators blinded to treatment groups.

In the same cohort used for gut histology, circulating levels of the bacterial endotoxin LPS and the proinflammatory cytokine IL-17A were determined using established ELISA assays (Earnest et al., 2022). Serum LPS levels were quantitatively measured using a double-sandwich ELISA method (MyBiosource, USA) and colorimetric detection system. Serum and standard samples (100μl/well) were assayed in duplicate. Assay sensitivity ranged from 15.6-1000ng/ml. Plates were read at 450nm in a plate reader (BioTek), and sample measurements were interpolated from the standard curve. Serum levels of the cytokine IL-17A were measured in blood samples from fixed and shifted LD animals using a multiplexed magnetic bead immunoassay (Milipore Corp. MA) and a Bio-Plex suspension array system (Bio-Rad Laboratories, CA) following manufacturer’s protocols.

### Statistical analysis

Statistical analysis was performed on SCFA, IL-17A and LPS data to determine the significance of LD treatment (fixed versus shifted) and time (pre-versus post-treatment) differences using a two-way ANOVA adjusted for multiple comparisons in conjunction with Sidák’s post-hoc pairwise analysis. For gut morphology analysis, LD treatment group differences were evaluated using an unpaired *t*-test. Pearson’s correlation coefficients were determined to analyze the relationship between the abundance of beneficial gut bacteria and stroke survival. In each case, LD treatment- and/or time-related differences in circulating levels of SCFA, IL-17A and LPS and in gut morphological features were considered significant at p<0.05 (GraphPad, San Diego, CA).

## RESULTS

Stable entrainment of the circadian rhythm of wheel-running activity was observed in all animals during baseline exposure to standard LD 12:12 conditions. Consistent with the results of our previous studies (Earnest et al. 2016, Earnest et al. 2022), activity rhythms remained stably entrained with daily onsets of activity occurring shortly after lights-off in fixed LD rats but were severely desynchronized in the shifted LD group throughout the period of exposure to experimental lighting conditions (see Earnest et al., 2016).

16S Amplicon sequencing analysis yielded important information on treatment group differences in microbiome composition, albeit limited to the genus level and relative quantification with minimal functional characterization. Alpha diversity metrics revealed decreased richness (Observed ASVs metrics, **Fig. 2A**) as well as evenness (Shannon Index, **Fig. 2B**) in male rats in the shifted LD group (pre vs post: p<0.001, and p<0.001, respectively) but not in fixed controls (p=0.052, and 0.4, respectively). Female rats, instead, showed no significant differences in richness (**Fig. 2D**) in the shifted (p=0.3), or fixed LD groups (p=0.8). However, female rats had a significantly higher Shannon index (**Fig. 2F**) in the shifted LD group (pre vs post: p=0.001) but not in fixed LD controls (p=0.8).

**Figure 2:**
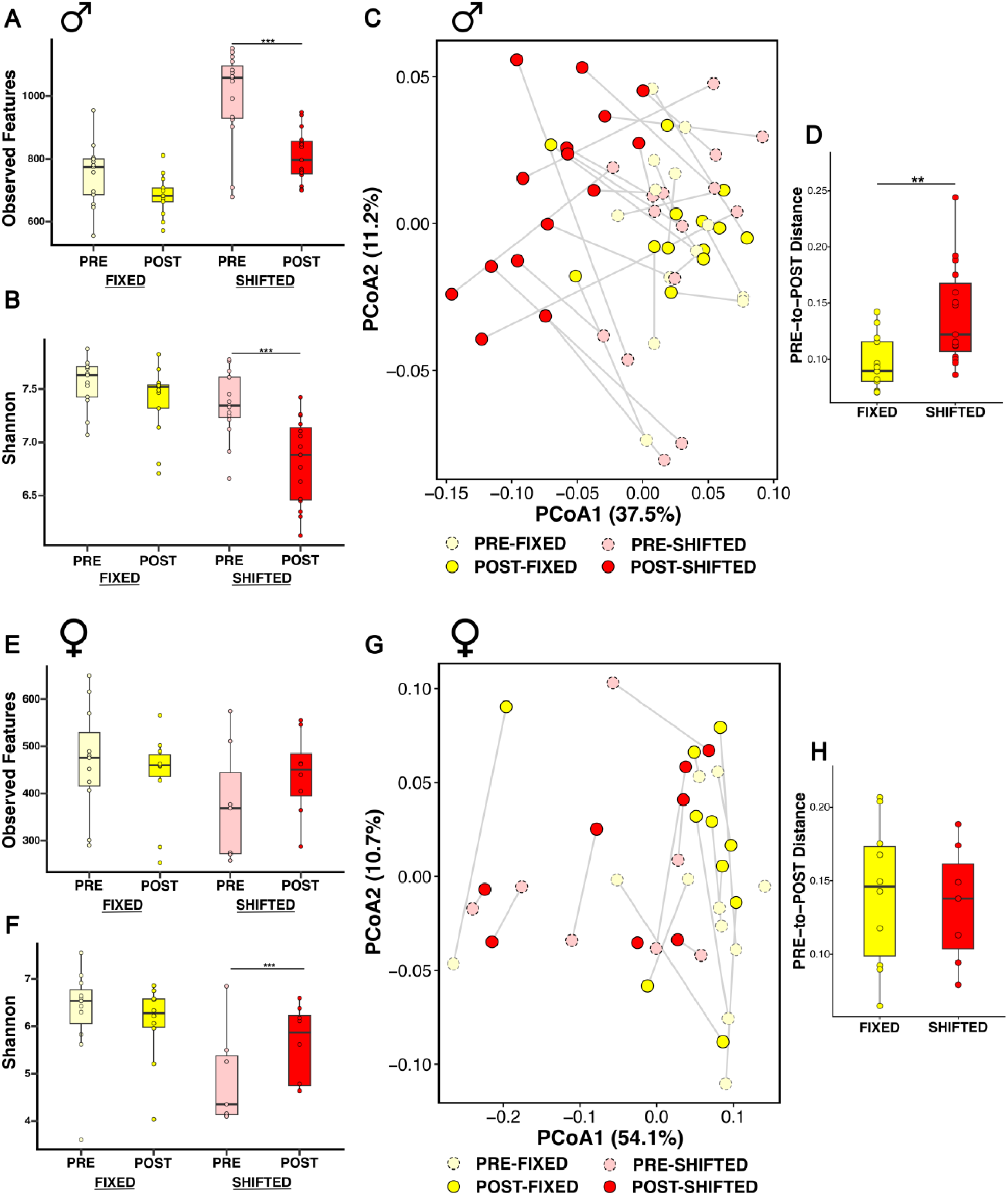
Effects of shifted LD cycles on microbiome composition. Microbial community composition in males and females maintained under fixed or shifted LD cycles. Alpha diversity, assessed as richness (Observed Features; **A**, **E**) and evenness (Shannon index; **B**, **F**), was measured at pre- and post-timepoints for each condition in males (**A**, **B**) and females (**E**, **F**). Beta diversity was calculated using weighted UniFrac principal coordinate analysis (PCoA) to assess changes in overall community structure between timepoints for males (C) and females (**G**). Weighted UniFrac boxplots of intra-individual difference in pre vs. post distances for males (**D**) and females (**H**) exposed to fixed or shifted LD cycles. ***, p<0.001; ** p<0.01.

Beta diversity in male rats, calculated through the weighted UniFrac distance matrix, showed significant differences in overall microbial community composition within both shifted and fixed LD groups over time (PERMANOVA, p=0.001 and 0.032, respectively). However, the portion of variance related to time was different, with a larger effect observed in the shifted LD group (R²=0.29), whereas fixed LD male rats showed high overlap (R²=0.08), both of which are depicted graphically in **Fig**. **2C**. Alternatively, no significant differences were observed in fixed or shifted LD female rats over time (PERMANOVA, p=0.1, R²=0.08; p=0.1, R²=0.05), as reflected by the high degree of overlap in the PCoA plots (**Fig. 2G**). Intra-individual differences between pre- and post-distances had significantly larger change over time in shifted LD males compared to their fixed counterparts (p=0.003, **Fig. 2D**). No significant change in intra-individual differences between pre- and post-distances was observed between shifted and fixed LD females (p=0.7, **Fig. 2H**).

Differential abundance analysis with MaAslin3 at the genus level identified several taxa that were differentially abundant over time within fixed and shifted LD rats. Genera characterized by significant associations with the combined effects of time and circadian dysregulation are shown in Tables 1 (male rats) and 2 (female rats). In males, pre-versus post-comparisons revealed 8 significantly associated taxa in fixed LD controls and 24 in shifted LD rats at significance level q-value joint <0.05. Of interest, post-treatment abundance of SCFA-producing taxa such as *Harryflintia* and *Prevotellaceae (UCG-001, UCG-003)* were significantly reduced in shifted LD male rats when compared to baseline levels (**Table 1**). Moreover, the significant decreases in post-treatment abundance of *Butyricimonas*, *Bacteroides*, and *Rikenellaceae* in shifted LD male rats are notable because some members of these taxa are considered potential pathobionts that may become pathogenic or promote inflammation under certain conditions (Chow, Tang, and Mazmanian, 2011), Further analysis using Pearson correlations revealed significant positive correlations between the abundance of several beneficial gut bacteria and stroke survival (**Fig. 3**). In both shifted and fixed LD males, a positive relationship was observed between stroke survival (i.e., number of days that animals survived after stroke) and the relative abundance of *Akkermansia* (p<0.01, r= 0.7308), *Butyricicoccus* (p<0.01, r= 0.6810), and *Prevotella* (p<0.01, r= 0.7289). The positive correlation between the abundance of these beneficial bacteria and stroke survival is noteworthy in relation to the reported functions of *Akkermansia* in maintaining gut barrier integrity, and inhibiting intestinal inflammation by suppressing the production of inflammatory cytokines (Reunanen et al., 2015; Pan, Barua and Ip, 2022; Mo et al., 2024), *Butyricicoccus* in producing the neuroprotective SCFA butyrate (Russell et al., 2013; Cavaliere et al., 2022), and *Prevotella* in modulating neuroinflammation (Houlden et al., 2016; Khatoon et al., 2023; Zhuang et al., 2025). In contrast, females had only 1 association with significant pre-versus post-difference in fixed LD controls and 6 in the shifted LD group. In accord with our findings that mortality was low in females and that stroke survival (in days) did not differ between treatment groups (Earnest et al. 2016), no significant positive or negative correlations were observed between the abundance of these beneficial bacteria and stroke survival in shifted or fixed LD female rats.

**Figure 3:**
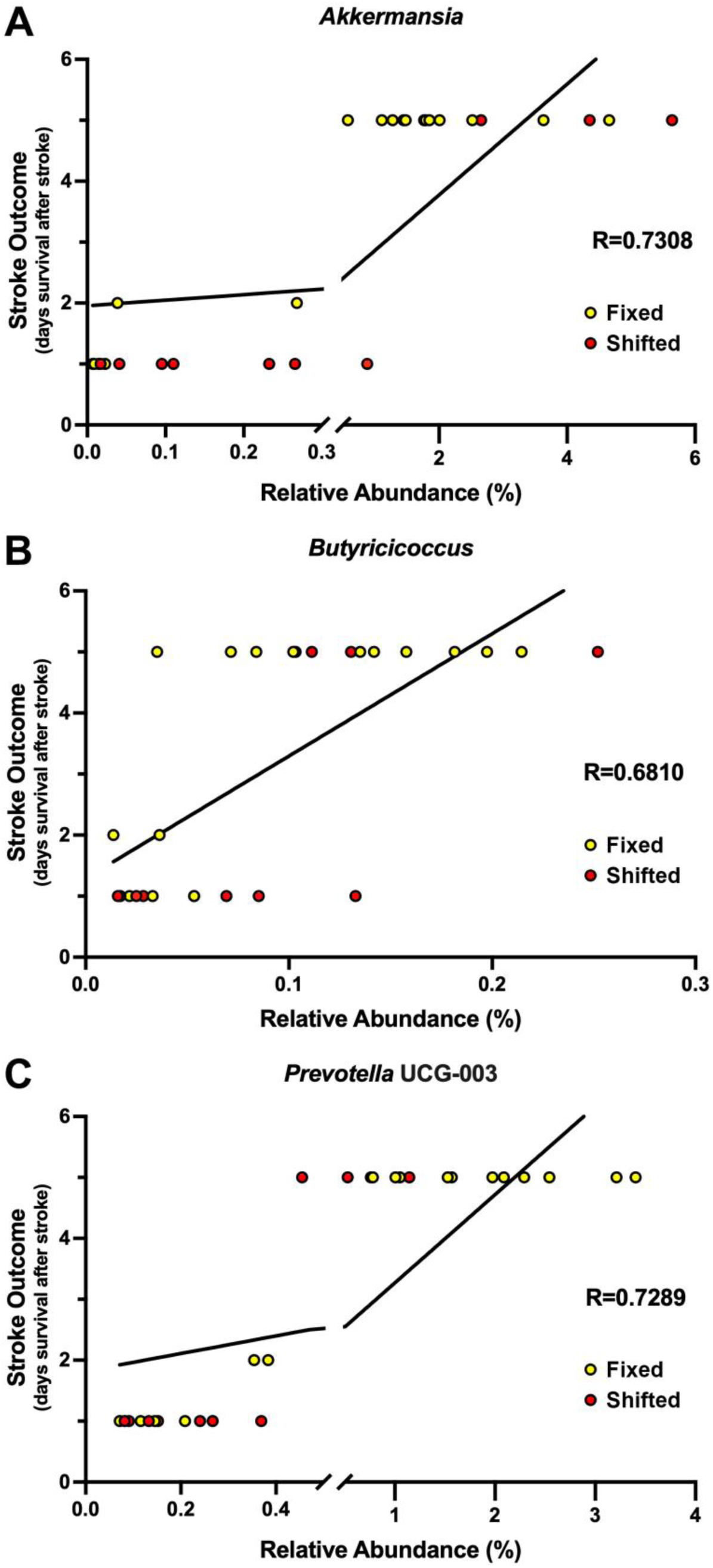
Relationship between the fecal abundance of beneficial gut bacteria and stroke survival in male rats that were exposed to fixed or shifted LD cycles. Pearson correlation coefficients comparing stroke survival (days after ischemic stroke) with the relative abundance of: (A) *Akkermansia*, (B) *Butyricicoccus*, and (C) *Prevotella* in cohorts of male rats exposed to fixed (n=12-13) or shifted (n=15) LD cycles. Symbols represent individual data values for each rat in fixed (yellow) and shifted (red) LD groups. Lines in each graph denote simple linear regression for the data set with corresponding *p* values.

**Table 1.**
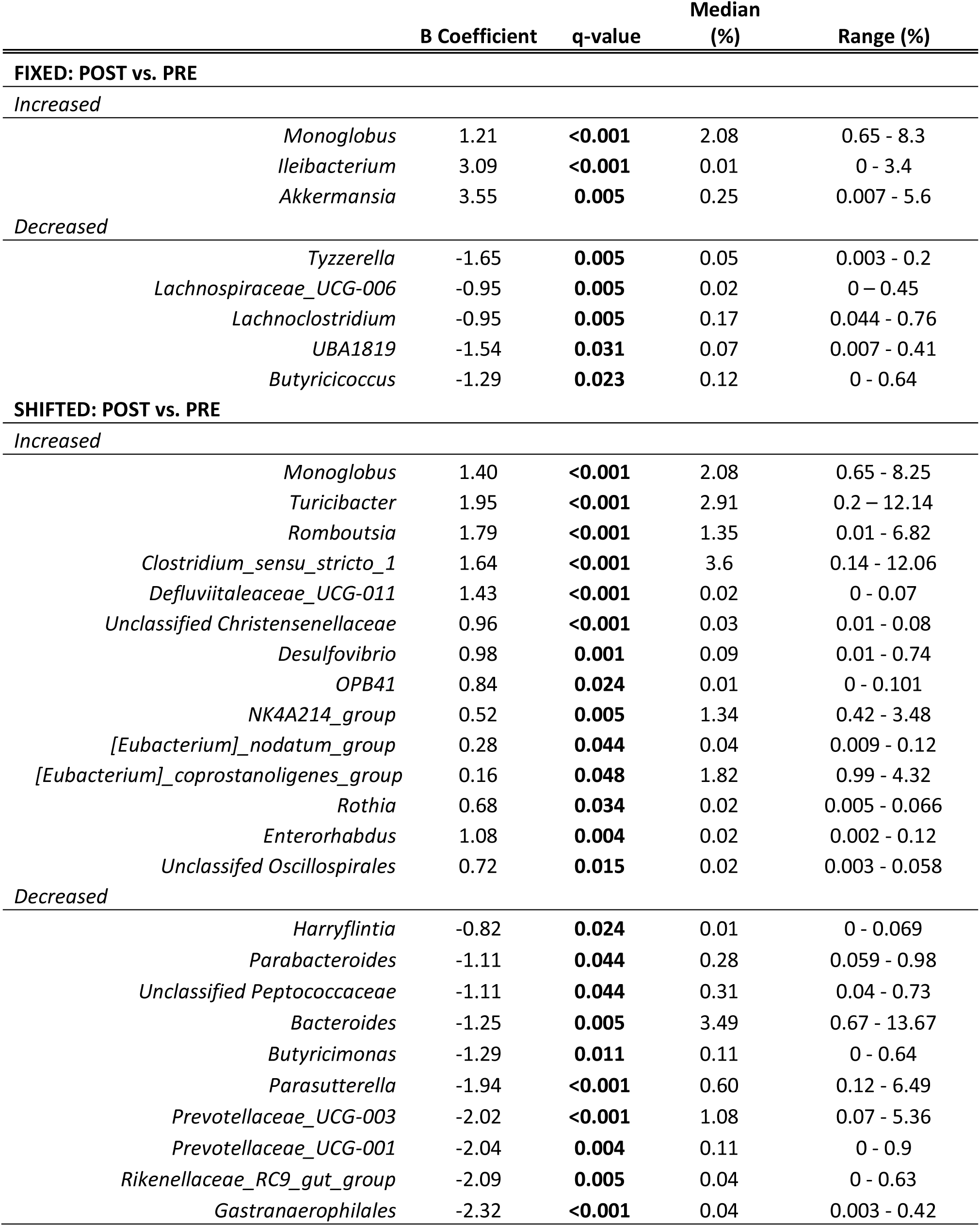
MaAsLin3 differential abundance statistics for males.

**Table 2.** MaAsLin3 differential abundance statistics for females.

| | Genus | $\beta$ Coefficient | q-value | Median (%) | Range (%) |
| --- | --- | --- | --- | --- | --- |
| **FIXED: POST vs. PRE** |  |  |  |  |  |
| *Increased* |  |  |  |  |  |
|  | *Frisingicoccus* | 3.32 | **0.049** | 0 | 0 – 0.10 |
| *Decreased* |  |  |  |  |  |
|  | *Paraprevotella* | -3.00 | **0.043** | 0 | 0 – 0.24 |
|  | *Prevotellaceae_UCG-003* | -2.33 | **0.046** | 0.05 | 0.004 – 3.35 |
| **SHIFTED: POST vs. PRE** |  |  |  |  |  |
| *Decreased* |  |  |  |  |  |
|  | *Unclassified Coriobacteriales* | -0.93 | **0.002** | 0.39 | 0.009 – 0.17 |
|  | *Adlercreutzia* | -0.41 | **0.005** | 0.26 | 0.073 – 0.85 |
|  | *Enterorhabdus* | -0.71 | **<0.001** | 0.17 | 0.016 – 0.75 |
|  | *Unclassified Ruminococcaceae* | -1.31 | **<0.001** | 0.5 | 0.016 – 4.61 |

Gut microbes critically affect neural function through the production of metabolites such as SCFAs (Alkasir et al., 2017), which serve a variety of beneficial roles including the modulation of immune cell activation and maintenance of the blood-brain barrier and blood-gut barrier. Because butyrate is a well-known SCFA with demonstrated roles as an anti-inflammatory mediator and neuroprotectant in response to stroke (Russell et al., 2013, Park and Sohrabji, 2016), we examined whether the effects of circadian dysregulation on gut microbiome composition are coupled with a decrease in the extravasation of this beneficial gut metabolite. In both sexes, circulating levels of total SCFAs were not significantly different over time (pre- vs post-treatment) in either fixed or shifted LD rats (**Fig. 4**). In contrast, sex differences were manifested in the effects of circadian rhythm dysregulation on serum levels of butyrate. Following experimental LD treatments, serum butyrate levels in male rats were significantly decreased in the shifted LD group relative to baseline pre-treatment values (p<0.01) but showed no significant differences in fixed LD controls when compared to basal levels (p=0.5004). In females, post-treatment levels of butyrate were not significantly different in fixed (p=0.5446) or shifted (p=0.4206) LD rats when compared to baseline levels in both groups. Serum levels of the SCFAs, acetate, propionate, and valerate, in both sexes were not significantly altered over time (pre- vs post-treatment) in either fixed or shifted LD rats,

**Figure 4:**
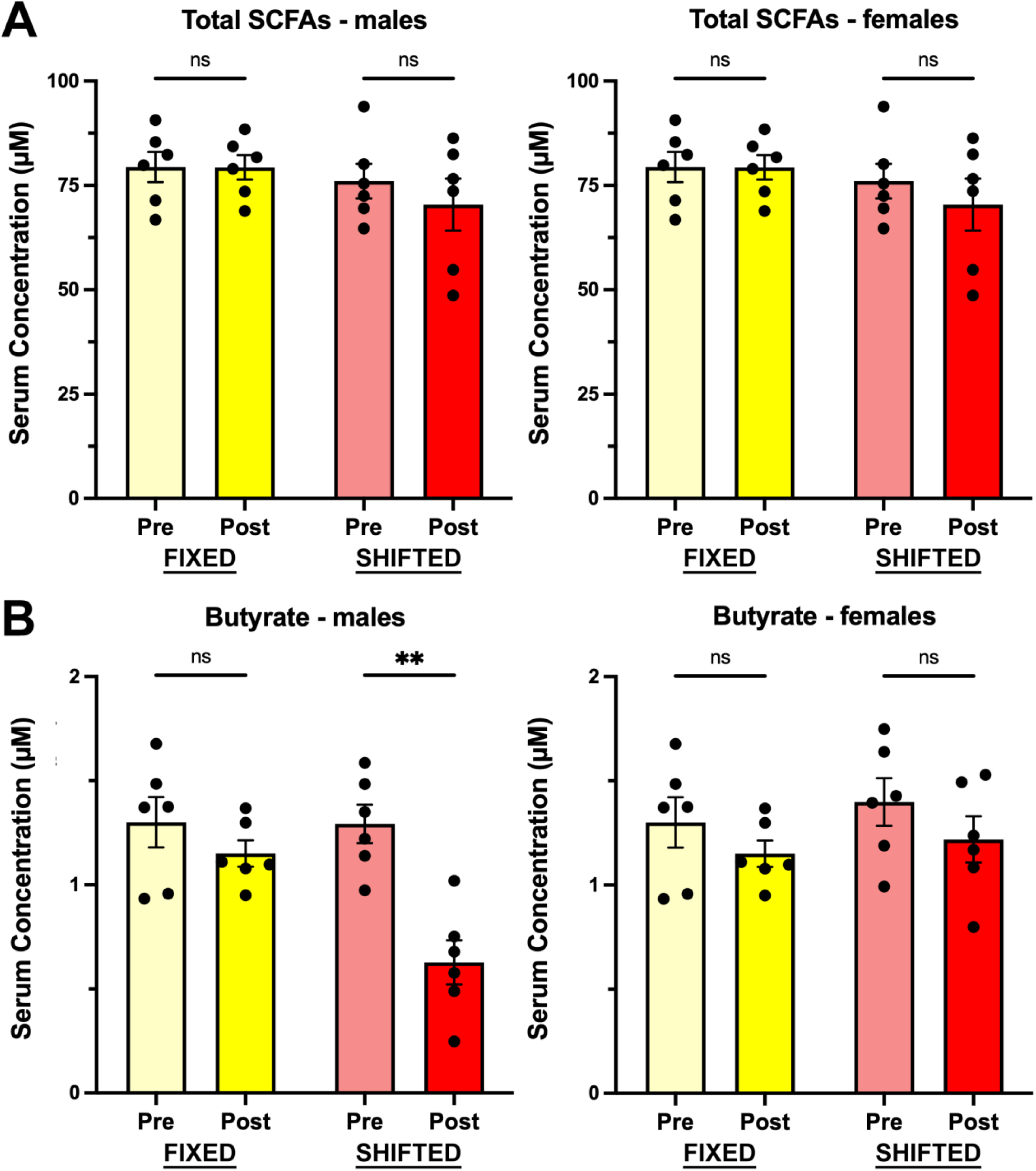
Effects of shifted LD cycles on circulating levels of short-chain fatty acids (SCFA). Serum levels of total SCFAs (**A**) and butyrate (**B**) in male (**left**) and female (**right**) rats exposed to fixed (yellow bars; n=6) or shifted (red bars; n=6) LD cycles. Histograms depict mean total SCFA and butyrate (µM) concentrations (+SEM) in blood collected immediately before (PRE) and after (POST) experimental LD cycle manipulations. *, p<0.05; **, p<0.01.

Histological analysis also revealed evidence of sex differences in the effects of circadian dysregulation on gut cytoarchitectural organization. In all fixed LD rats, gut morphology was characterized by the normally elongated and evenly spaced villi with a single row of crypt cells. In comparison, the distal ileum in shifted LD males, but not females, was distinguished by an “injured” appearance in which the villi were shorter, wider and blunted (**Fig. 5A)**. These decreases in villus length and width in shifted LD males effectively serve to reduce the villi surface area, thereby lowering nutrient absorption in the ileum. Due to the reductions in both villus length and width, male rats exposed to shifted LD cycles were correspondingly marked by a significant decrease (p<0.05) in the villus length/width ratio relative to their male counterparts in the fixed LD group (**Fig 5B**).

**Figure 5:**
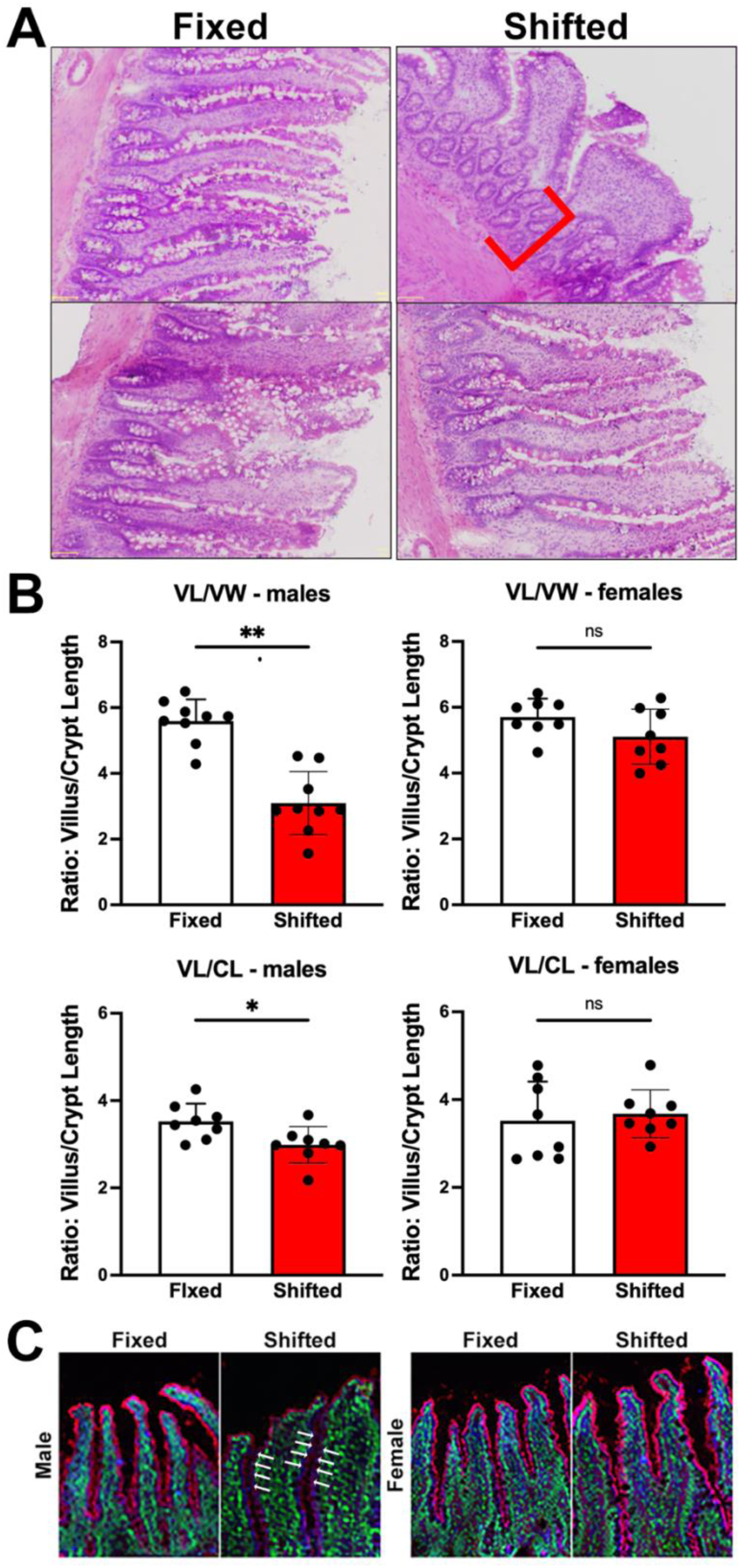
Effects of shifted LD cycles on gut morphology and barrier integrity. (**A**) H&E-stained sections of the ileum from male (**top**) and female (**bottom**) rats exposed to fixed (**left**) or shifted (**right**) LD cycles. Crypt hyperplasia and increased crypt length (red bracket) were evident in shifted LD males. (**B**) Histograms depict mean (± SEM) ratio of villus length:width (**top**) and villus:crypt length (**bottom**) for each group (n=8). (**C**) Immunohistochemistry for the tight junction protein ZO-1 (red) and villin (green) in sections (10µM) of the distal ileum from male and female in the fixed and shifted LD groups. White arrows indicate the location of the epithelial barrier where inter-epithelial ZO-1 fluroescence was virtually absent in shifted LD males. *, p<0.05; **, p<0.01.

In addition to shorter, blunted villi, the “injured” gut appearance in shifted LD males was accompanied by crypt hyperplasia. Crypt cells in the intestine provide stem cells that aid in repair of the epithelium as well as growth and protection of the ileum. Crypt hyperplasia was differentially observed in male rats exposed to shifted LD cycles (**Fig. 5B**), resulting in a significant decrease (p<0.05) in villus to crypt length ratio as compared to fixed LD controls of the same sex. This crypt hyperplasia and corresponding reduction in villus to crypt length ratio are indicative of increased pro-inflammatory responses in the gut and the impaired function of tight junction proteins to maintain gut barrier integrity. Thus, circadian dysregulation had marked effects on gut morphology and barrier integrity in males, but not females, exposed to shifted LD cycles including deceases in villus length:width ratio along with crypt hyperplasia and disruption of the gut epithelial barrier.

The integrity of the gut barrier was further examined using the pattern of immunohistochemical localization of the tight junction protein, ZO-1 and villin, a cytoskeletal protein expressed in the brush border of epithelial cells lining the gastrointestinal tract. Similar to the observed gut dysmorphology in response to circadian dysregulation, this analysis demonstrates that the gut barrier epithelium was disrupted exclusively in male rats exposed to shifted LD cycles. ZO-1 immunofluorescence was localized contiguously along brush border of the villi in fixed LD males as well as in females from both treatment groups (**Fig. 5C**). However, the gut barrier was distinctly altered in shifted LD male rats such that ZO-1 immunostaining along the brush border was absent or greatly diminished.

Because circadian rhythm dysregulation promotes a chronic basal inflammatory state (13, 14), we next examined the effects of the shifted LD paradigm on the endotoxin LPS and cytokine IL-17A, which act synergistically to intensify proinflammatory signaling and activate immune cells (Waisman et al., 2015, Kawanokuchi et al., 2008). In both fixed and shifted LD rats, serum levels of LPS (**Fig. 6A**) and IL-17A (**Fig. 6B**) were analyzed at baseline (pre-treatment) and immediately after exposure to experimental lighting conditions (post-treatment). In accord with previous reports (El-Hakim et al., 2021), sex differences were observed in circulating levels of bacterial endotoxin LPS such this gut-derived inflammatory mediator was much higher in males, irrespective of treatment (fixed, shifted) and timing (pre-vs post-treatment). In both fixed and shifted LD males, post-treatment LPS levels were elevated over time (F_(1,26)_: 18.58; p=0.00020). Planned post-hoc comparisons revealed that after experimental LD treatments serum LPS levels in male rats were significantly elevated in the shifted LD group relative to baseline pre-treatment values (p<0.01) but showed no significant differences in fixed LD controls when compared to basal levels (p=0.6569). In females, a significant interaction effect was observed (F_(1,28)_:17.14; p=0.0003) between the pre- and post-treatment levels of LPS, and planned comparisons indicate that post-treatment levels of serum endotoxin were significantly increased (p<0.05) in the shifted LD group but were significantly decreased (p<0.05) in fixed LD controls when compared to baseline levels in both groups.

**Figure 6:**
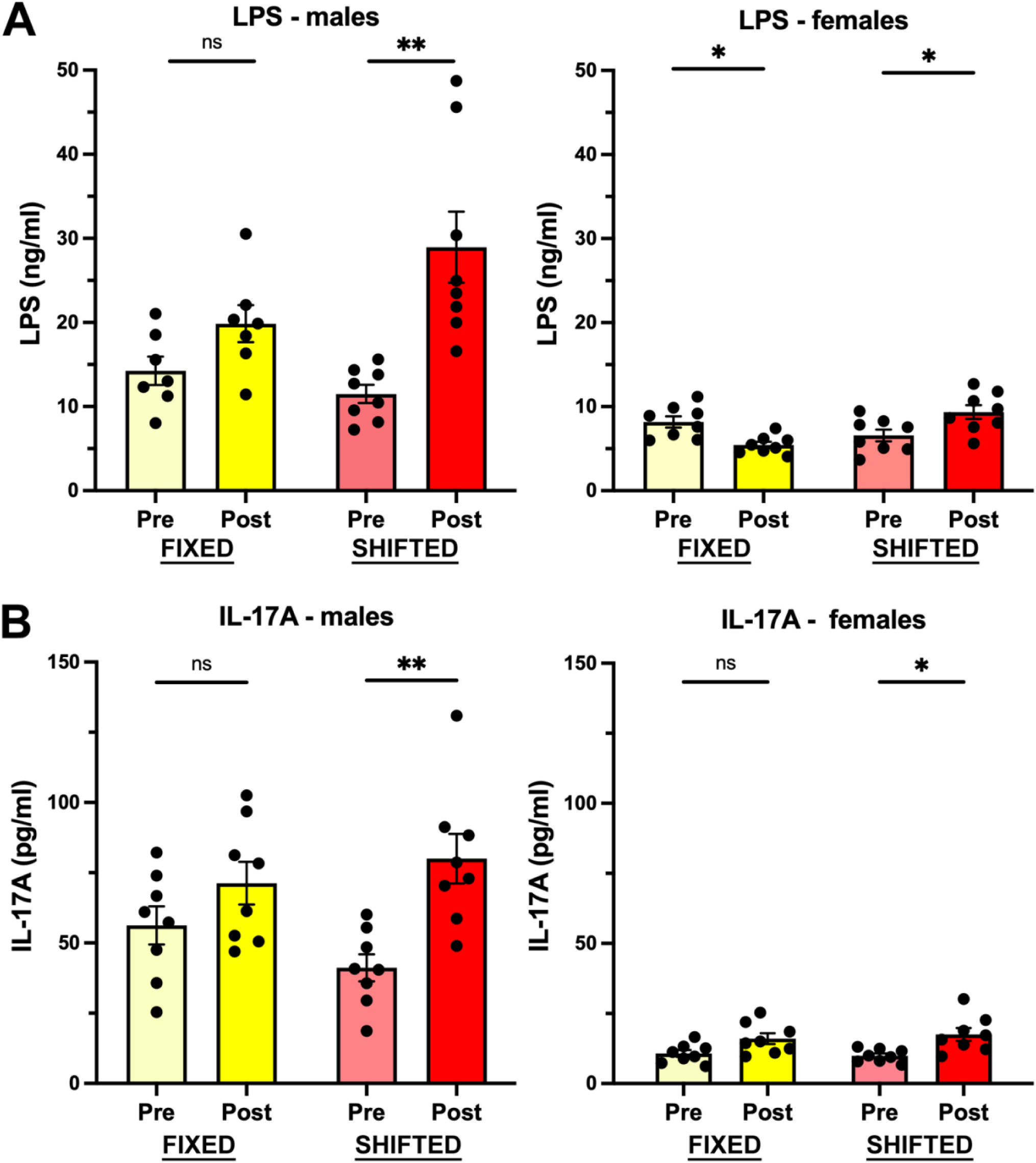
Effects of shifted LD cycles on circulating levels of lipopolysaccharide (LPS) and the inflammatory cytokine IL-17A. Serum levels of LPS (**A**) and IL-17A (**B**) in male (**left**) and female (**right**) rats exposed to fixed (yellow bars; n=7-8) or shifted (red bars; n=8) LD cycles. Histograms depict mean LPS (ng/ml) and IL-17A (pg/ml) levels (+SEM) in blood collected immediately before (PRE) and after (POST) experimental LD cycle manipulations. *, p<0.05; **, p<0.01.

Circulating levels of IL-17A in male rats were elevated in both the LD fixed and shifted groups over time compared to their respective baseline values (F_(1,28)_:14.02; p=0.0008) (**Fig. 6B**). However, post-treatment IL-17A levels were significantly increased (p<0.01) by approximately 2-fold in the shifted LD males but were not significantly different (p=0.6169) in fixed LD controls of the same sex relative to baseline levels observed in both groups. In females, a similar effect was observed in which IL-17A was elevated in the serum of both treatment groups over time (F_(1,28)_:15.10; p=0.0006) and post-treatment levels of this inflammatory cytokine were significantly increased in both fixed (p<0.05) and shifted (p<0.01) LD rats relative to baseline values in each group.

## DISCUSSION

Circadian dysregulation and the extent of stroke outcomes have been independently associated with alterations in the gut microbiome and barrier function. Under normal conditions, gut contents are partitioned from general circulation via a series of barrier elements, including the intestinal epithelial barrier and the blood-gut barrier (BGB) (26). Increased permeability of the BGB not only allows transfer of gut contents but also alters the composition of resident microbes, and consequently, synthesis of beneficial gut metabolites. Disruption of circadian rhythms during exposure to constant light has been shown to increase intestinal permeability, decrease diversity of gut microbiota and dramatically downregulate beneficial butyrate-producing microbial communities (Deaver et al., 2018). In turn, gut barrier breakdown and bacterial translocation are associated with more severe stroke outcomes (Stanley et al., 2016; Park et al., 2020). The present study provides further evidence that circadian rhythm dysregulation and stroke severity are closely interconnected through changes in gut health. It is noteworthy that sex-differences were manifested in the impact of circadian dysregulation on the gut microbiome. Male rats exposed to shifted LD cycles were distinguished by decreased alpha diversity, altered beta diversity, and significant reductions in beneficial bacterial taxa such as *Harryflintia* and *Prevotellaceae*. In contrast, female rats showed only modest microbiome alterations and no significant disruptions in overall diversity metrics. This sex divergence closely mirrors epidemiological and experimental data indicating that circadian dysregulation interacts with biological sex to influence disease susceptibility (Karlsson et al., 2005; Earnest et al., 2016). The observed reduction of SCFA-producing genera in males is particularly relevant, given the neuroprotective roles of SCFAs in maintaining blood–brain barrier and gut barrier integrity, regulating immune responses, and mitigating stroke-induced injury (Russell et al., 2013; Park & Sohrabji, 2016).

Exposure to shifted LD cycles produced pronounced disruptions in intestinal cytoarchitecture and barrier integrity in male, but not female, rats. Histological and immunohistochemical analyses revealed that the distal ileum in shifted LD males was marked by shortened and blunted villi, crypt hyperplasia, and disrupted or absent localization of the tight junction protein ZO-1, indicative of impaired absorptive capacity and increased epithelial permeability (’leaky gut’). Importantly, the absence of these morphological alterations in females exposed to shifted LD cycles demonstrates a sex-specific vulnerability of the male gut epithelium to circadian dysregulation. Consistent with this compromise of gut barrier integrity, serum endotoxin (LPS) concentrations were markedly elevated in shifted LD males, whereas females exhibited only modest post-treatment changes in LPS levels. In parallel, circulating levels of IL-17A were substantially increased relative to baseline in shifted LD males but exhibited much smaller elevations in shifted LD females, IL-17A, a cytokine produced primarily by Th17 and γδ T cells, has been shown to promote crypt cell proliferation and epithelial remodeling (Waisman et al., 2015; Zepp et al., 2017), providing a mechanistic basis for the observed crypt hyperplasia in males exposed to shifted LD cycles. Moreover, IL-17A synergizes with LPS to amplify proinflammatory cascades within both the gut and central nervous system (Sun et al., 2015; Zhang et al., 2005). These findings are consistent with published reports on sex differences in the effects of changes in the gut microbiome and barrier integrity on stroke outcomes (Durgan et al., 2019; Ahnstedt et al., 2020; El-Hakim et al., 2021; Honarpisheh, Bryan and McCullough, 2022) and collectively support a model in which circadian dysregulation initiates a feed-forward loop in males: epithelial barrier disruption facilitates endotoxin leakage, which in turn promotes IL-17A upregulation, further exacerbating barrier dysfunction and driving a persistent systemic inflammatory state. This male-biased response of the gut–immune axis may provide a vital link between circadian dysregulation and exaggerated injury after ischemic stroke, whereas the relative preservation of gut architecture, microbiome composition and attenuated inflammatory responses in females may underlie their observed resilience to the pathological impact of circadian-related disturbances.

Taken together, our findings suggest that circadian rhythm dysregulation amplifies stroke severity through a gut–brain axis mechanism involving changes in gut microbiome composition and barrier function, altered SCFA metabolism, and systemic inflammation. Importantly, these changes are more pronounced in shifted LD males, aligning with our prior observations of higher stroke mortality in circadian-disrupted males compared to females (Earnest et al., 2016). Beyond stroke, the convergence of circadian misalignment, gut dysfunction, and inflammation has broad relevance to human health, as shift work has been linked to elevated risk for cardiovascular disease, diabetes, obesity, cancer, and Alzheimer’s disease-related dementias (Torquati et al., 2018). Our data provide mechanistic support for these associations, implicating sex-specific gut pathophysiology as a potential driver.

While this study identifies compelling sex differences in gut and immune responses to circadian dysregulation, several limitations warrant consideration. First, although we observed correlations between microbial taxa abundance and stroke survival, causality cannot be inferred. Future studies employing microbiome transfer or targeted microbial supplementation will be critical to establish whether restoring beneficial taxa (e.g., *Akkermansia*, *Butyricicoccus*, *Prevotella*) can mitigate circadian dysregulation-induced stroke severity. Second, we focused on the ileum, but circadian rhythm dysregulation may differentially affect other intestinal segments. In addition, we acknowledge that exercise may have some implications in the sex-dependent effects of circadian rhythm dysregulation on stroke outcomes because: 1) exercise associated with running wheel activity improves stroke outcomes (Marin et al., 2003; Hayes et al., 2008); and 2) sex-, but not, treatment-dependent differences in the total amount of daily wheel-running behavior were observed in our previous stroke study (Earnest et al., 2016) such that daily activity levels were approximately 10-fold greater in female subjects than in their male counterparts within each treatment group. Finally, although our model simulates rotating shift work, it does not capture lifestyle variables such as diet, stress, or socioeconomic factors that often accompany to circadian misalignment in individuals where workplace or social influences commonly impose highly irregular schedules on sleep-wake patterns, mealtimes and other health-related processes.

In summary, this study demonstrates that controlled circadian dysregulation differentially perturbs gut architecture, microbiome composition, metabolite extravasation, and inflammatory signaling in a sex-dependent manner, with males showing greater vulnerability. These gut-mediated changes provide a plausible mechanism by which circadian dysregulation exacerbates stroke outcomes and may also contribute to other chronic diseases linked to shift work. The gut–brain axis thus represents a critical target for interventions aimed at mitigating the adverse health effects of circadian misalignment.

